# Comparative root transcriptome analysis suggests down-regulation of nitrogen assimilation in DJ123, a highly phosphorus-efficient rice genotype

**DOI:** 10.1101/2024.04.18.590184

**Authors:** M. Asaduzzaman Prodhan, Yoshiaki Ueda, Matthias Wissuwa

**Affiliations:** School of Biological Sciences, The University of Western Australia, 35 Stirling Highway, Perth, WA 6009, Australia; Crop, Livestock and Environment Division, Japan International Research Center for Agricultural Sciences (JIRCAS), Ohwashi 1-1, Tsukuba, Ibaraki, Japan

**Keywords:** DJ123, IR64, Phosphorus, Nitrate, Ammonium, Transcriptome

## Abstract

Many cultivable lands across the globe are characteristically low for plant-available phosphorus (P). This necessitates application of P fertilisers, but this increases farming costs beyond the affordability of marginal farmers. Thus, developing cultivars with high P-use efficiency (PUE) is necessary in high-yielding modern rice varieties, which are typically inefficient in P usage. However, the molecular and physiological bases to increase PUE in crops remain elusive. Here, we studied root transcriptomes of two breeding parents contrasting in PUE via RNA-seq to elucidate key physiological and molecular mechanisms that underlies efficient use of P in rice. Examination of transcriptome data obtained from plants grown under P-sufficient and P-deficient hydroponic conditions in DJ123 (an upland rice genotype adapted to low P soils) and IR64 (a modern rice variety less efficient in P use) revealed that the genes encoding nitrogen assimilation-related enzymes such as glutamine synthetase [EC. 6.3.1.2], glutamate synthase [EC. 1.4.1.13], and asparagine synthetase [EC. 6.3.5.4] were down-regulated only in DJ123 roots while it was not significantly affected in IR64 under low P conditions. In addition, DJ123 roots had a lower total nitrogen (N) concentration than IR64 irrespective of P conditions. Taken together, we surmise that the low level of N concentration together with down-regulation of the N assimilation-related genes allow DJ123 to operate at a low level of N, thus leading to formation of root tissues with lower metabolic investment and a greater PUE.

## Introduction

Rice is staple food for more than half of the world population^1^. The demand for rice production is ever increasing to feed the growing population^2^. However, total rice cultivable lands are shrinking due to urbanisation and industrialisation^3,4^, and rice yields are adversely affected by numerous biotic and abiotic stresses, among which improper supply of soil nutrients is one of the major issues. Among the essential nutrients for plants, rice fields are characteristically low in plant-available phosphorus (P)^5,6^, and supplementation of P-fertilisers is necessary to obtain maximum yield. However, farmers in developing countries, which often coincide with P-deficient soils, often cannot afford sufficient P fertilisers, and rice plants in these areas are severely affected due to the lack of P-efficient high yielding rice varieties^7^. In light of continuously increasing P-fertiliser price, developing P-efficient rice varieties is indispensable to meet the growing global rice demand^8^.

P-efficient rice variety will have the properties like high P-acquisition efficiency (PAE) and/or P-use efficiency (PUE)^9^. PAE is an efficiency that enhances P acquisition from the rhizosphere. The traits that lead to a greater PAE will include longer roots, distribution of roots on shallower layer of the soil, more vigorous root branching, denser root hairs, reduced root diameter, mycorrhizal associations etc^6,10,11^. On the other hand, PUE is an internal P-efficiency that maximises the usage of the internal P by producing more yield and biomass per unit P^12^. Consequently, properties associated with high PUE will ultimately reduce the demand for external P fertilisers while achieving higher yield and biomass.

There are some rice landraces that are low-yielding but possess remarkable P-efficiency traits. For example, an *aus*-type landrace originating from south Asia, DJ123^13^, has the genetic potential that allows greater PAE^14^ and PUE^15^, thus it performs exceptionally well in low P soils^16^. Therefore, there is a global breeding effort to transfer the PAE and PUE properties from the low-yielding P-efficient genotypes to the modern high yielding P-inefficient genotypes^17^. For example, IR64 is a widely cultivated rice variety because of its higher yields under optimal conditions^18^. However, IR64 loses its yield potential in low P soils because of its P-inefficiency^19^.

To accelerate the breeding efforts, it is crucial to understand how DJ123 attains such a higher PAE and PUE in low P soils. The genetic mechanisms that orchestrate a greater PAE and PUE in DJ123 are still unknown. A recent transcriptomic study suggested that DJ123 changes its global gene expression patterns faster or more strongly upon a short period (24 h) of P deficiency in root as compared with IR64, based on the number of differentially expressed genes^20^. It suggests that certain molecular mechanisms allow DJ123 to respond to external low P status more sensitively than the modern-cultivar IR64, which could partly account for high PAE and PUE. On the other hand, the effects of persisting low P availability on global gene expression patterns have not been compared in DJ123 and IR64. We anticipate that a comparative transcriptomic analysis might reveal clues of these underlying mechanisms that enable DJ123 sustain vigorous growth under low P conditions.

Here, we compared the root transcriptomes of P-efficient DJ123 under a long-term low-P condition with those of P-inefficient IR64. Cultivation of each individual plant in separate bottles allowed each genotype to absorb exactly the same amount of P, allowing a precise evaluation of PUE, since PUE negatively correlates with PAE. Through investigation of differentially expressed genes and their functional annotation, we suggest that low P-induced down-regulation of genes involved in nitrogen (N) metabolic pathway as a candidate mechanism for high PUE in DJ123.

## Materials and methods

### Plant materials and growth conditions

Rice (*Oryza sativa* L.) genotypes DJ123 and IR64 were grown hydroponically in a naturally lit, temperature-controlled glass house at Japan International Research Center for Agricultural Sciences, Tsukuba, Japan; at a mean temperature of 27 °C (23–33 °C) and a mean relative humidity of 40% (10– 78%) between September and November 2019. Plants were grown in 1.1-L light-impermeable bottles containing modified Yoshida solution for 42 days as described previously^20^. At the end of the treatment period, roots were harvested, snap-frozen immediately in liquid nitrogen and stored at − 80 °C until RNA extraction.

### Determination of total N concentration

Total nitrogen (N) content were measured using 41-d-old plants deriving from our previous report^21^, which were grown in the 1.1-L light-impermeable bottles under the same environment with nutrient solution containing low or high concentrations of P using a mass spectrometry (Delta V Advantage, Thermo Fischer Scientific).

### RNA sequencing

TRIzol reagent (Invitrogen, Carlsbad, CA, USA) was used to extract total RNA from the fine-ground root materials following manufacturer’s instructions. Extracted RNA was treated with DNase and quantified using a NanoDrop spectrophotometer (Thermo Scientific). The integrity was checked using an Agilent 2100 Bioanalyzer (Agilent Technologies, Inc., Santa Clara, CA, USA). RNA samples with a RIN value ranging from 6.2 to 8.6 were subjected to sequencing library preparation using TruSeq Stranded mRNA Low Throughput (LT) Sample Preparation Kit (Illumina, San Diego, CA, USA). The prepared library was sequenced using NovaSeq6000 sequencer (Illumina) to generate 150-bp paired-end reads.

The quality of the sequencing reads was checked using FastQC (http://www.bioinformatics.babraham.ac.uk/projects/fastqc/). The raw reads were trimmed using trimmomatic software^22^ using the following criteria; i) trimming the Illumina PE adapters (TruSeq3-PE-2.fa), ii) removal of N stretches from the ends that had a quality score below 3, iii) discarding the poor quality sequences that had a quality score below 15 in a 4-base sliding window, and iv) exclusion of the reads with less than 25 bases in length. The quality filtered RNA-seq reads were mapped to the rice reference genome (*Oryza sativa* Japonica Group, https://plants.ensembl.org/info/data/ftp/index.html) using HISAT2^23^. The alignment files were converted to ‘sam’ files^24^ and subjected to StringTie^25^ for transcript assembly.

### Differential gene expression analysis

We quantified the transcripts for differential gene expression analysis using DESeq2^26^. We applied a Generalized Linear Model (design = ∼ Genotypes + Treatments + Genotypes:Treatments) for differential gene expression analysis and discarded genes that had less than 10 counts. Genes with ‘False Discovery Rate’ threshold at 0.005; padj at < 0.001; and fold change range by < 0.5 or >2.

### Gene Ontology (GO) term enrichment analysis

The differentially expressed genes were passed on to GO term analysis under a series of ‘Annotation Data Set’ viz. ‘GO biological process complete’, ‘GO molecular function complete’, ‘GO cellular component complete’, ‘PANTHER Pathways’, ‘PANTHER GO-Slim Biological Process’, ‘PANTHER GO-Slim Molecular Function’, ‘PANTHER GO-Slim Cellular Component’ and ‘PANTHER Protein Class’ that are implemented in PANTHER classification system (http://pantherdb.org/tools/compareToRefList.jsp)^27^. For the annotation, ‘Fisher’s Exact’ test with ‘No correction’ was applied against the ‘*Oryza sativa* all genes in the database’ as a reference list.

### Validation of the RNA-seq data

The results of RNA-seq were validated by quantitative real-time PCR (qRT-PCR) assay of selected genes using a CFX96 Touch Real-Time PCR system (BioRad, USA) as described previously^20^.

## Results

### Transcriptomes show phosphorus-dependent genotypic variation

To investigate genotypic differences in responsive pattern to low P, we performed a comparative root transcriptome analysis using P-efficient DJ123 as a highly P use-efficient genotype under low P condition and IR64 as a P inefficient modern rice cultivar and grew them under P deficient conditions for 41 days after germination. The PCA analysis showed that the genotypes explained 67% of the variation, while 18% was explained by P treatments (Fig. 1A). Following DEG analysis revealed that low P condition induced differential expression of 2,258 and 2,156 genes in DJ123 and IR64 respectively (Fig. 1B). These genes contained 1,250 up-regulated genes and 1,008 down-regulated genes in DJ123 (Fig. 1C), and 1,228 up-regulated and 928 down-regulated genes in IR64 (Fig. 1D). Among these DEGs, only 894 were common in both genotypes, while the rest were genotype specific (Fig. 1B).

**Figure 1.**
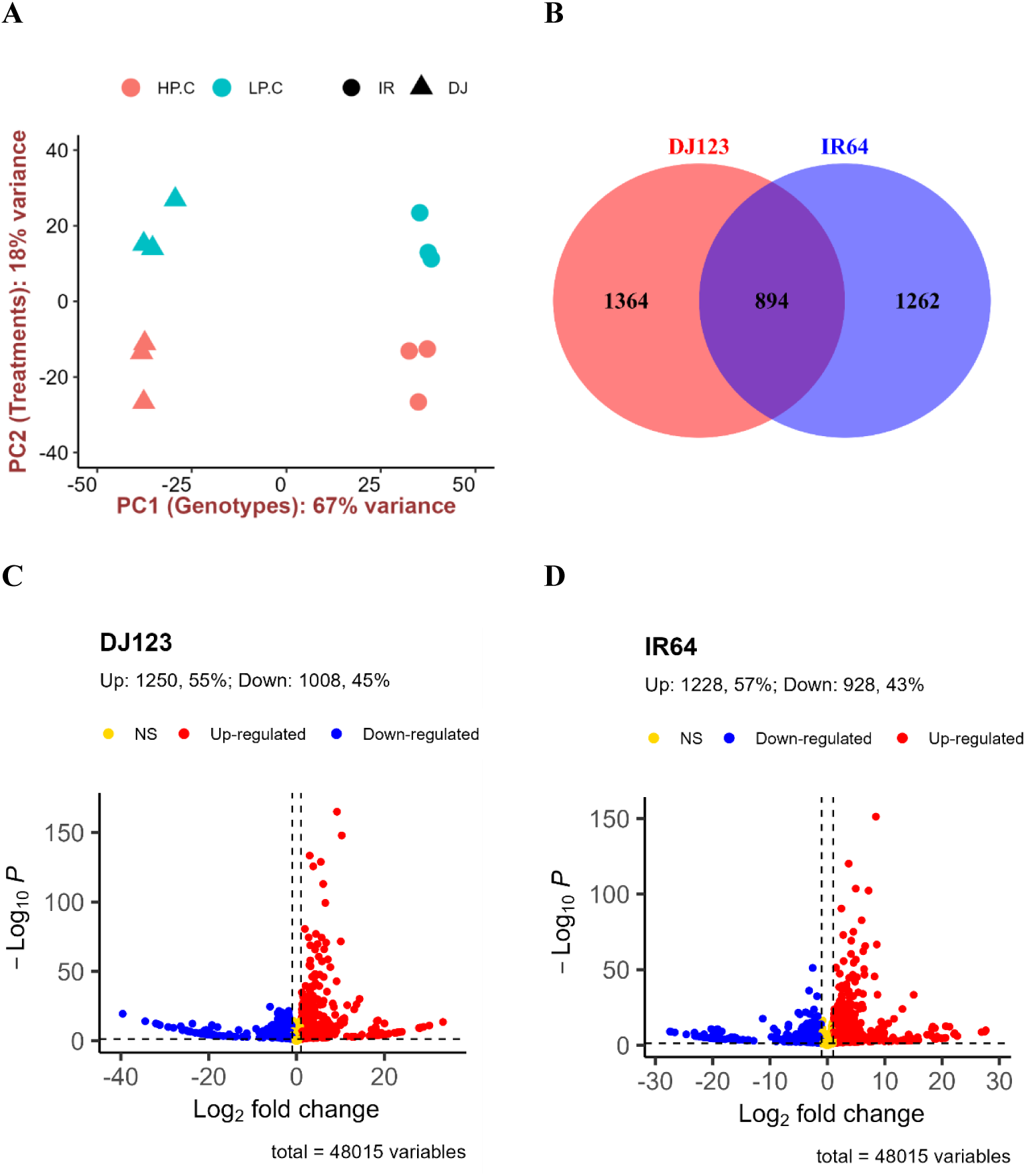
Summary of RNA-seq results. **A)** Principal Component Analysis of the RNA-seq results. HP and LP indicates “high P” and “low P”. **B)** Venn diagram of the differentially expressed genes (DEGs) in roots of DJ123 and IR64. The number of DEGs in LP treatment against HP treatment is shown. **C, D)** Volcano plots showing significantly up- or down-regulated genes in DJ123 **(C)** and IR64 **(D)** roots by LP treatment. The logarithm (log_2_) of fold change of each transcript is presented on the x-axis, and -log_10_(*p*-value) is on the y-axis. In **B-D**, thresholds for DEG were ‘adjusted *p*-value < 0.05’ and ‘log_2_(Fold Change) > 1’.

### Down-regulated genes in DJ123 are enriched with N assimilation-related Gene Ontology terms

We carried out gene ontology (GO) term analysis to functionally annotate the group of genes induced or suppressed by low P treatment in DJ123 and IR64 roots (Figs. 2-3, Suppl. Figs. 1-6). Genes induced by low P treatment were enriched with Molecular Function GO terms such as ‘inorganic phosphate transmembrane transporter activity’, and ‘glycerophosphodiester phosphodiesterase activity’ that were common in both genotypes (Suppl Fig. 3). Protein Class GO Terms like ‘transporter’, ‘transferase’, ‘phosphosdiesterase’, phosphatase, ‘metabolite interconversion enzyme’ and ‘hydrolase’ were also enriched in both DJ123 and IR64 by the low P induced genes (Suppl. Fig. 5). Interestingly, the down-regulated genes in DJ123 under low P condition were significantly enriched with the GO terms such as ‘nitrogen utilization’, ‘glutamine family amino acid metabolic process’, ‘glutamine family amino acid biosynthetic process’, ‘glutamate metabolic process’, ‘glutamate biosynthetic process’, and ‘ammonia assimilation cycle’ (Fig. 2) under the Biological process category; and the GO terms ‘glutamate synthase activity’, ‘glutamate synthase activity’, and ‘glutamate synthase (NADH) activity’ under the Molecular function category (Fig. 3). This was in contrast with IR64 (Figs. 2-3).

**Figure 2.**
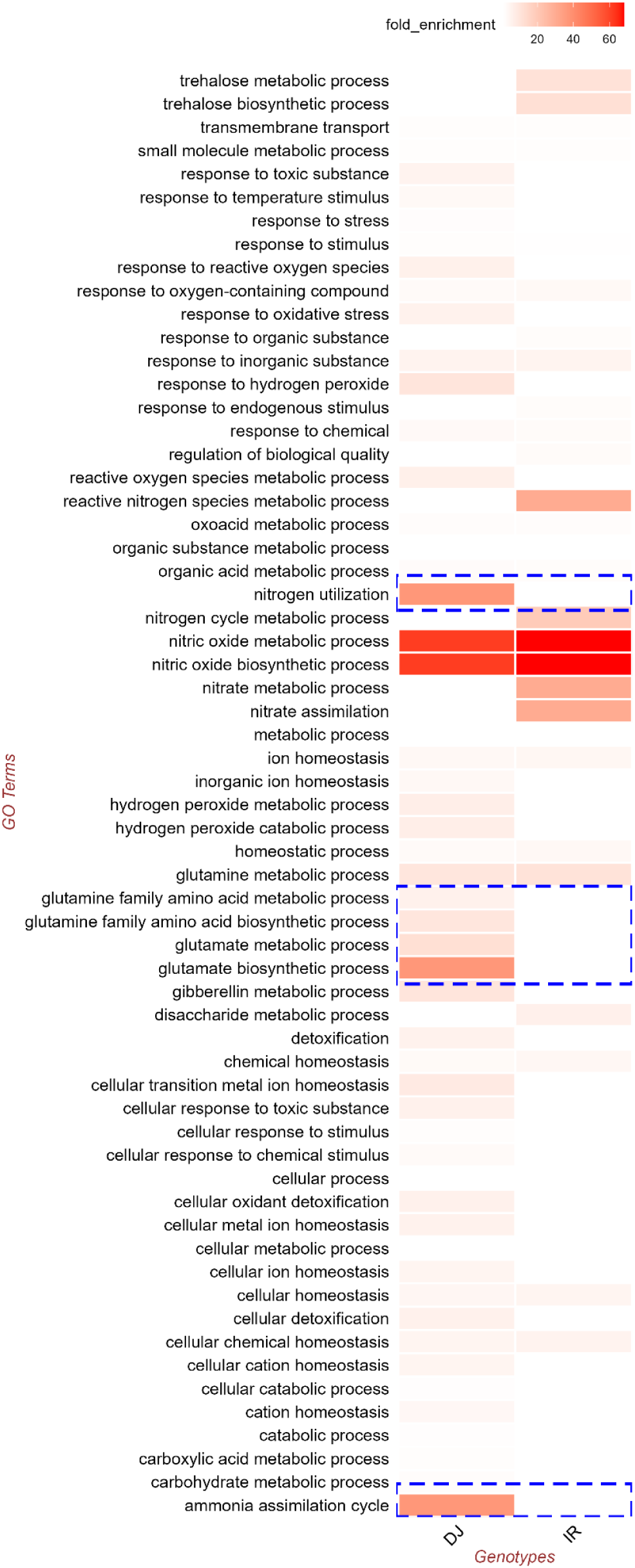
Biological processes (GO terms) enriched by the down-regulated genes in DJ123 and IR64 roots under low P treatment.

**Figure 3.**
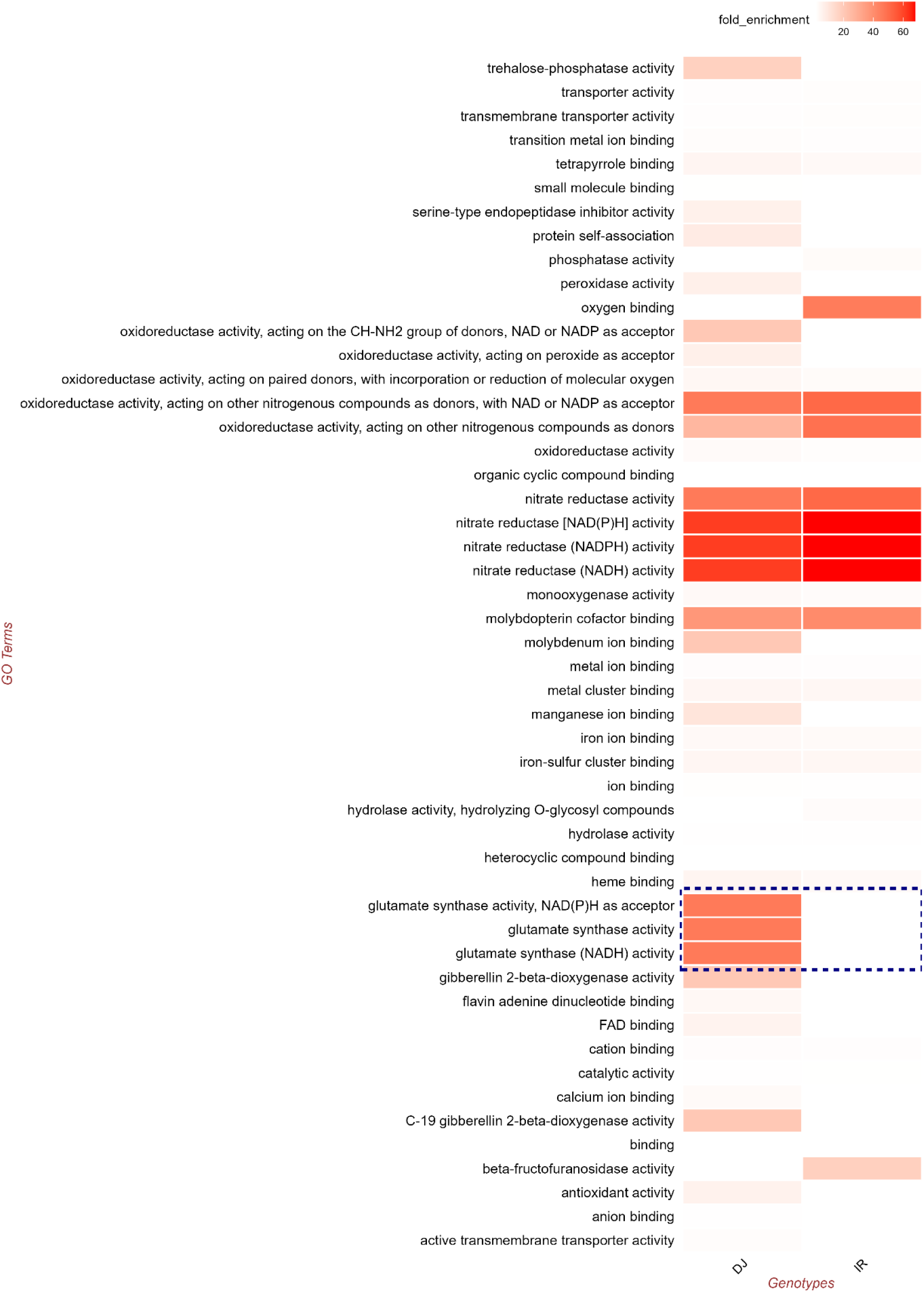
Molecular functions (GO terms) enriched by the down-regulated genes in DJ123 and IR64 roots under low P treatment.

### DJ123 down-regulated N assimilation-related genes under low P supply

We extracted the down-regulated genes in DJ123 that are associated with the GO terms specifically enriched in down-regulated genes in DJ123 under low P supply. According to The Rice Annotation Project Database (RAP-DB)^28,29^, these genes are annoted as pyrroline-5-carboxylate synthetase 2, glutamate synthetase 1, NADH-dependent glutamate synthase1, glutamate decarboxylase 3, NADH-glutamate synthase 2, glutamine synthetase 1;2, cytosolic glutamine synthethase 1;2, asparagine synthetase 1, and NADH-glutamate synthase 2 (Table 1). These genes are involved in the synthesis of Glu and Gln, the primary products of nitrogen assimilation via GS-GOGAT cycle (Yanagisawa, 2014), Glu metabolism and Asn synthesis (Fig. 4).

**Table 1.**
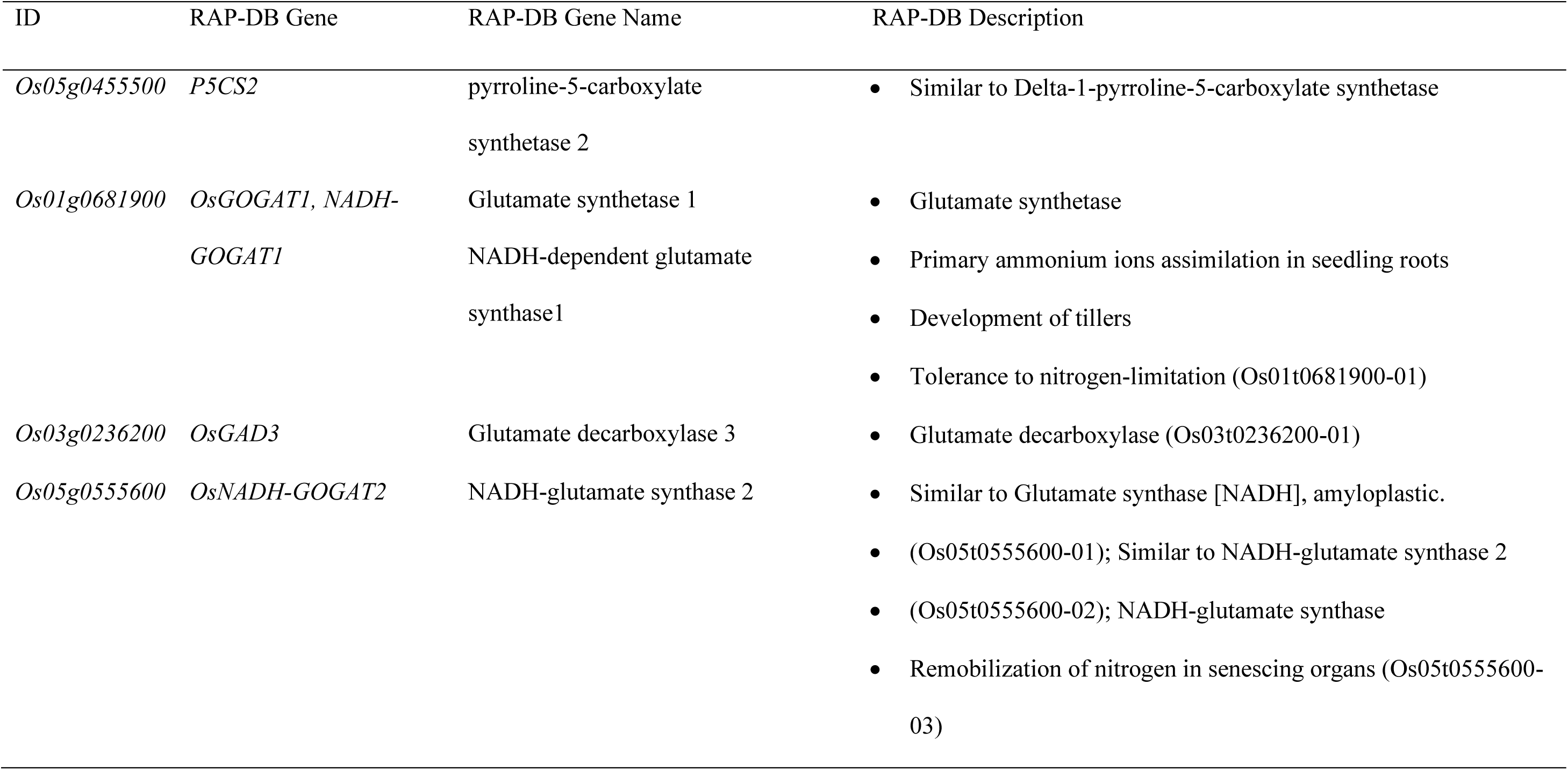
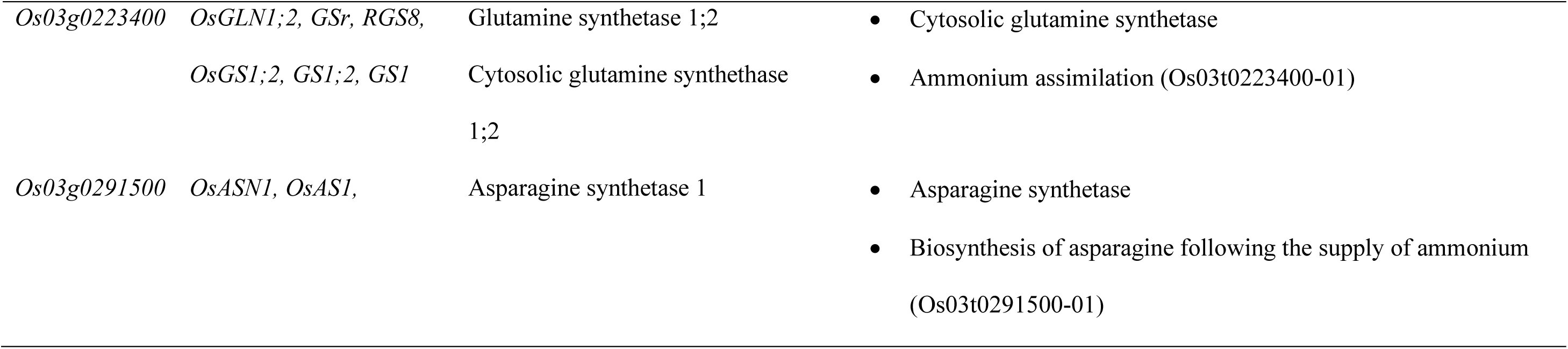
The down-regulated genes in DJ123 and IR64 roots under low P treatment mapped to the nitrogen assimilation and glutamine biosynthesis GO terms.

**Figure 4.**
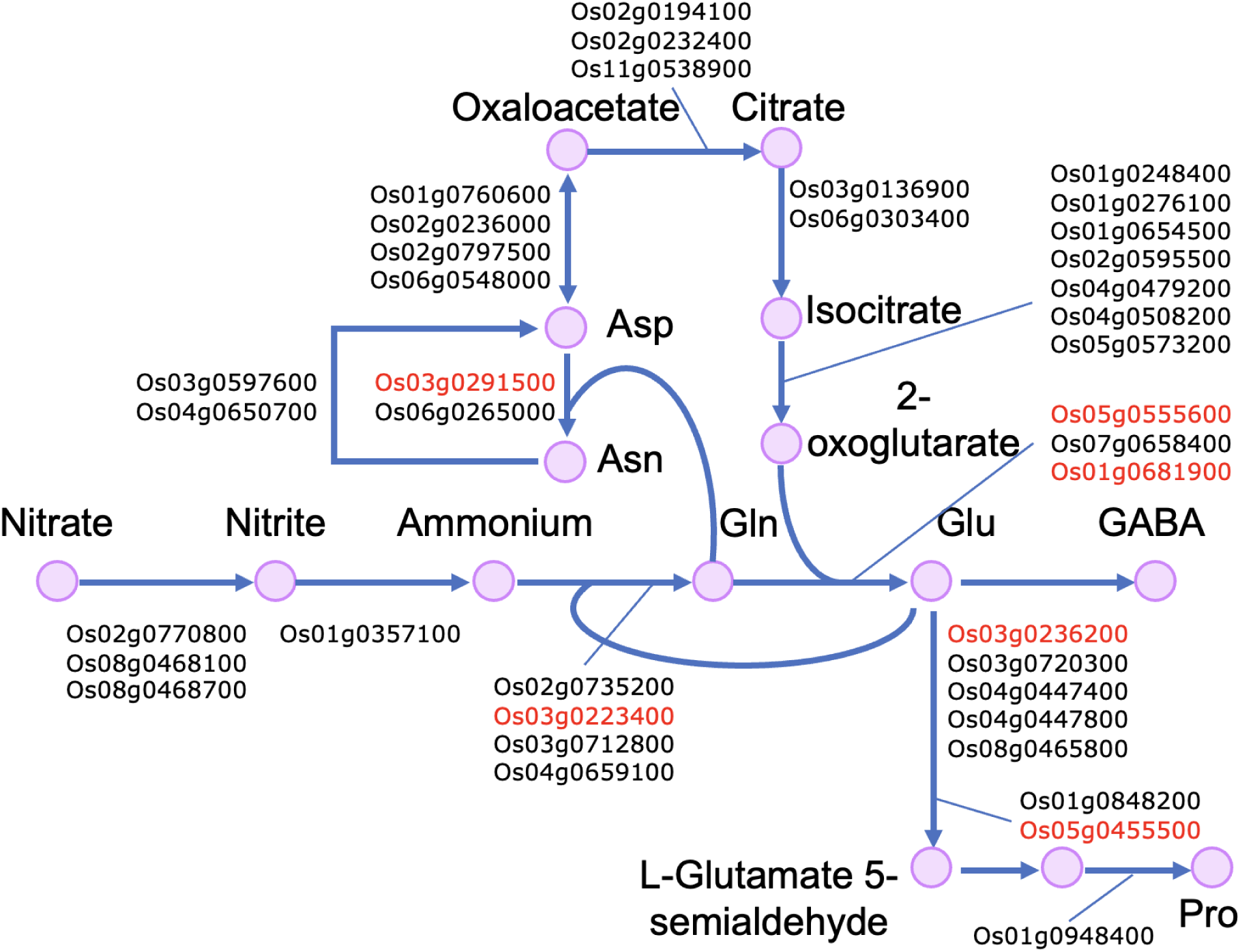
Metabolic pathways involved in N assimilation and some amino acids. Information on the metabolic pathway and associated genes was obtained from the KEGG database (ref). Gene IDs indicated in red are the genes significantly downregulated in DJ123 but not in IR64.

### DJ123 roots have lower total N concentration than IR64

Analysis of N content in DJ123 and IR64 plants in a separate experiment showed that roots of DJ123 had lower N concentrations than IR64 under low P supply (Fig. 5). Significant differences in total N concentrations was also observed under high P supply in root, while similar trend was not observed in shoot.

**Figure 5.**
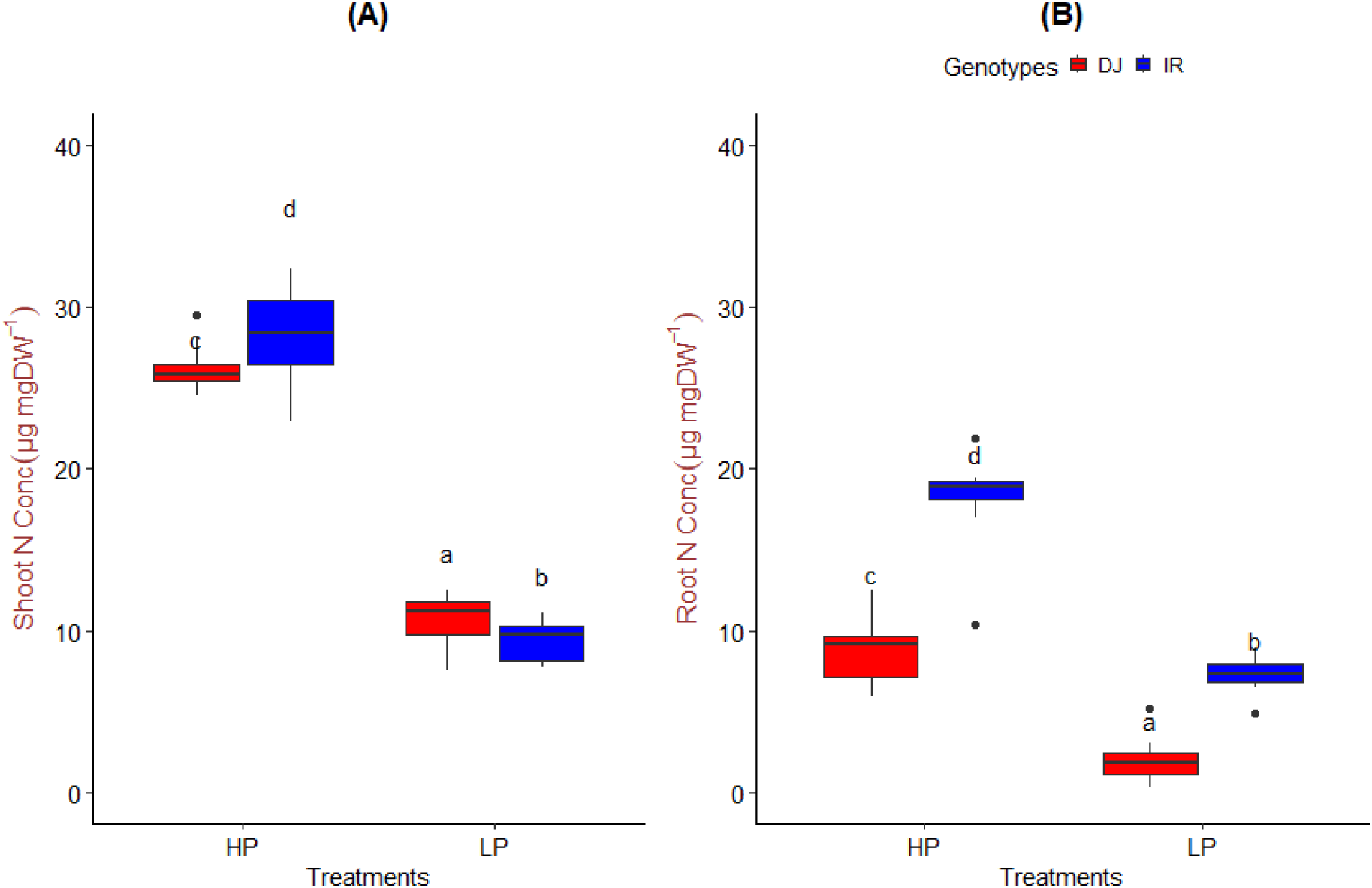
Total nitrogen (N) concentrations. Total N concentration in shoot **(A)** and root **(B)** of DJ123 and IR64 under high P (HP) and low P (LP) conditions. The boxplot shows the median value (n=8) by the horizontal line in the middle, 25th and 75th percentiles as box limits (interquartile range), minimum and maximum values as whiskers that are still within the range of 1.5 times the interquartile range. Data were statistically tested by the ‘generalized least squares’ (gls) model using ‘nlme’^44^ R package. The least-squares means were extracted from the above gls model using the ‘emmeans’ function of the ‘emmans’^45^ R package. Then, multiple pairwise comparison of the least-squares means were carried out using the ‘multicomp::cld’ function of the ‘multicomp’^46^ R package adjusted for ‘Sidak’^47^ correction. Significant differences (*P* < 0.05) between the genotypes in each tissue are indicated by different alphabets.

## Discussion

Our PCA and differentially expressed genes (DEGs) analysis in roots of DJ123 and IR64 demonstrated that there was a genotypic difference between DJ123 and IR64 in responding to low P. Our previous study suggests that DJ123 responds to a short-term P deprivation more strongly than IR64^20^. On the contrary, under a long-term low-P treatment, both genotypes up- and down-regulated similar number of genes (Fig. 1C and D). However, only around 40% of genes were commonly affected by low P treatment in both genotypes (Fig. 1B), suggesting that P-efficient DJ123 and P-inefficient IR64 had distinctive response patterns to long-term P deficiency in root. Remarkably, GO term analysis of the genotype-specific DEGs revealed that genes involved in N assimilation were significantly downregulated in roots under low-P only in DJ123, while they were unchanged in IR64. These genes were i) *glutamine synthethase 1* and *2,* ii) *glutamate synthase 1* and *2*, iii) *pyrroline-5-carboxylate synthetase 2*, iv) *glutamate decarboxylase 3*, and v) *asparagine synthetase 1* (Fig. 4). Glutamine synthethase [EC. 6.3.1.2] is central player in ammonium assimilation in rice^30^. Glutamate synthase [EC. 1.4.1.13], which comprises the GS-GOGAT cycle together with glutamine synthetase, control ammonium assimilation in rice^31,32^. Pyrroline-5-carboxylate synthetase [EC. 2.7.2.11] catalises the first step of proline biosynthesis from glutamate^33^. Glutamate decarboxylase converts L-glutamate to γ-aminobutyric acid (GABA)^34^, which also serves as N storage molecule^35,36^. Despite its crucial role in N metabolism, a recent study reported that high GAD activity and resultant increase in GABA content confers tolerance to P deficiency in Arabidopsis. However, considering that the *GAD* homologue differentially regulated was downregulated in P-efficient genotype DJ123 in our current study, the contrasting regulation of *GAD* genes may not be the primary factor for P efficiency at least in roots in rice. Asparagine synthetase [EC. 6.3.5.4] converts glutamine to asparagine, an N storage amino acid^37^. Thus, down-regulation of these genes might suggest that DJ123 reduces the rate of N assimilation under low P supply. This was supported by the low level of nitrogen concentration in DJ123 roots under low P condition. However, considering that DJ123 roots also showed lower N concentration compared to that in IR64 under high P treatment, the molecular machineries that allow efficient use of P could be constitutively functional. It could be relevant with the fact that DJ123 is superior than IR64 to use other elements such as S compared with IR64.

Down-regulation of genes involved in N uptake or assimilation under P deficient conditions is considered to be an adaptive response, since the requirement of N decreases under P deficiency and circumventing excess N uptake and assimilation may save energy. Previous studies reported some mechanisms associated with down-regulation of N uptake and assimilation under P deficiency. In Arabidopsis, upregulation of a P deficiency-inducible transcriptional repressor NIGT1 suppresses the expression of a nitrate transporter *NRT2.1* and possibly some other N assimilation-related genes such as nitrate reductase and nitrite reductase^38^. Rice has only one *NIGT1* homologue, but its expression was not substantially different between high- and low-PUE genotypes nor between IR64 and DJ123 in root^21^, suggesting that the NIGT1-mediated regulation likely does not explain the contrasting regulatory patterns of N assimilation-related genes observed in the current study. Another possibility is repression of N-related genes by N-containing metabolites. It was reported that supplementation of several amino acids, especially Asn and Gln, followed by Glu and Ala, to roots of Arabidopsis significantly downregulated the expression of *NRT2.1* and nitrate uptake^39^. A previous metabolome study using leaves and roots of maize under P deficiency^40^ reported significant increases in amino acid contents in roots such as Glu and Ala. Taking into account that some genes involved in amino acid metabolism were differentially regulated in our current study, it is possible that genotypic differences in changes in amino acid profile affected the regulation of N-related genes between DJ123 and IR64. Further studies are necessary to validate if low rate of N assimilation is the key physiological factor for PUE and investigate underlying mechanisms for the contrasting regulation of N-related genes to get more profound insights into P use efficiency.

Considering together our results showing down-regulation of the N assimilation related genes and low N concentration in roots, we envisage that DJ123 might have genotypically evolved to operate at a low level of nitrogen regardless of P availability, which might orchestrate a greater PUE in low P soils. Our findings in rice corroborate with the Western Australian native plants that have evolved with tight control of nitrogen assimilation^41^ to adapt to the some of the most P-impoverished environments on earth^41–43^.

## Author contributions

M.A.P. and M.W. conceptualised the study. M.A.P. carried out the hydroponic experiment and RNA-seq, interpreted the analysis outputs and wrote the manuscript. Y.U. analysed the N content data. Y.U. and M.W. provided editorial comments. All authors critically discussed the results, read and approved the manuscript.

## Acknowledgements

We thank all the technical staffs at the Japan International Research Center for Agricultural Sciences (JIRCAS) for their technical assistance. We thank the Pawsey Supercomputer Centre (https://pawsey. org. au/) for the computing resources.

## Data availability statement

RNA-seq raw data analysed in this study were part of our RNA-seq project^20^ submitted to the National Centre for Biotechnology Information (NCBI) under BioProject ID PRJNA823747. This BioProject comprised of 72 samples. We have already published the results from 24 samples^20^. Here, we have analysed another set of 24 root samples treated with continuous ‘Low Phosphorus (LPC)’ and ‘High Phosphorus (HPC)’ for the entire experimental period. This manuscript reports the first and primary analysis of this dataset.

## Supporting information

**Supplementary Figure 1. Cellular Component (GO terms) enriched by the up-regulated genes in DJ123 and IR64 roots under low P treatment.**

**Supplementary Figure 2. Cellular Component (GO terms) enriched by the down-regulated genes in DJ123 and IR64 roots under low P treatment.**

**Supplementary Figure 3. Molecular Function (GO terms) enriched by the up-regulated genes in DJ123 and IR64 roots under low P treatment (Part1).**

**Supplementary Figure 4. Molecular Function (GO terms) enriched by the up-regulated genes in DJ123 and IR64 roots under low P treatment (Part2).**

**Supplementary Figure 5. Protein Class (GO terms) enriched by the up-regulated genes in DJ123 and IR64 roots under low P treatment.**

**Supplementary Figure 6. Protein Class (GO terms) enriched by the down-regulated genes in DJ123 and IR64 roots under low P treatment.**

