## Supplemental Material for "Comparative root transcriptome analysis suggests down-regulation of nitrogen assimilation in DJ123, a highly phosphorus-efficient rice genotype"

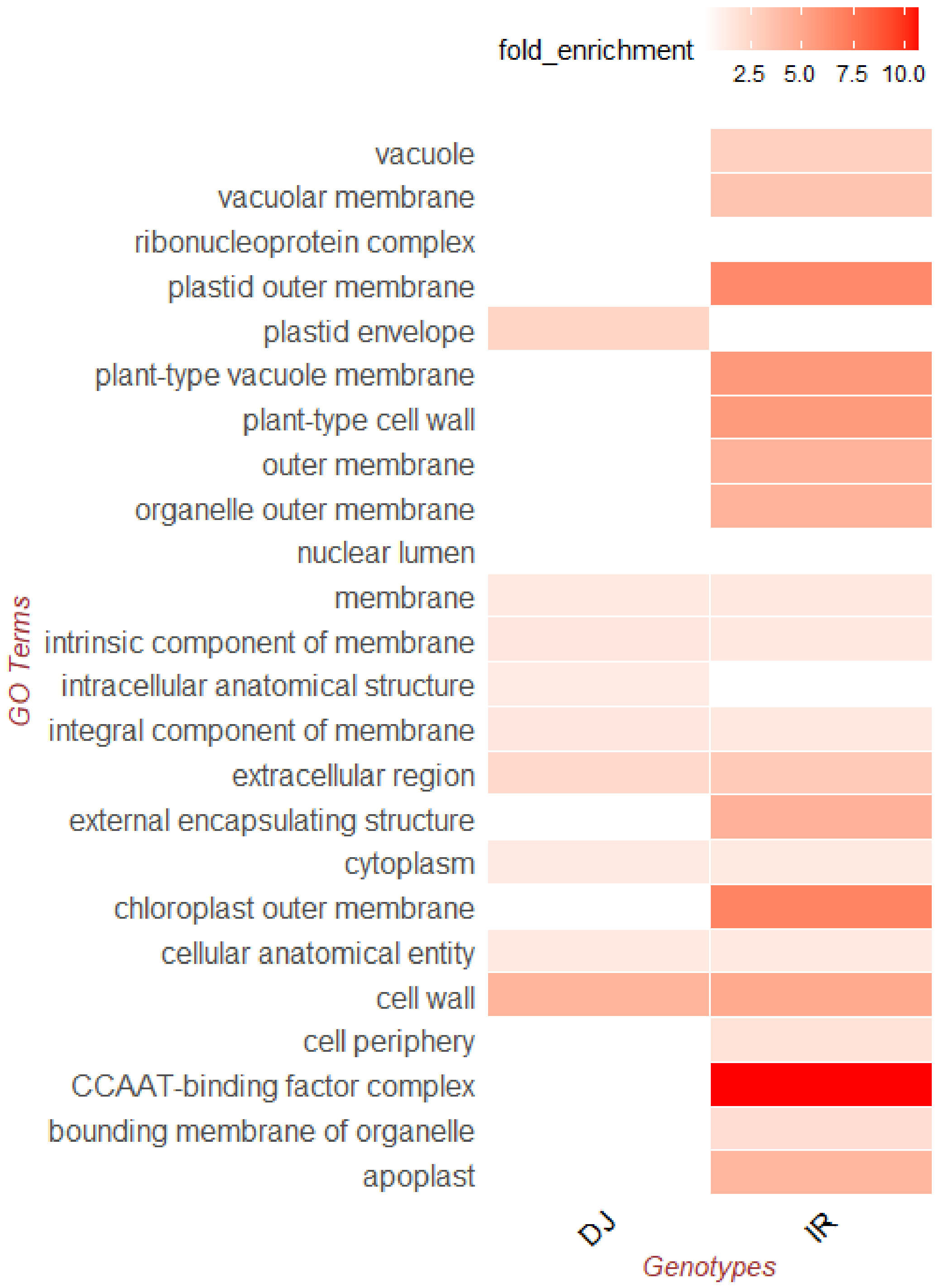


**Supplementary Figure 1.** Cellular Component (GO terms) enriched by the up-regulated genes in DJ123 and IR64 roots under low P treatment.

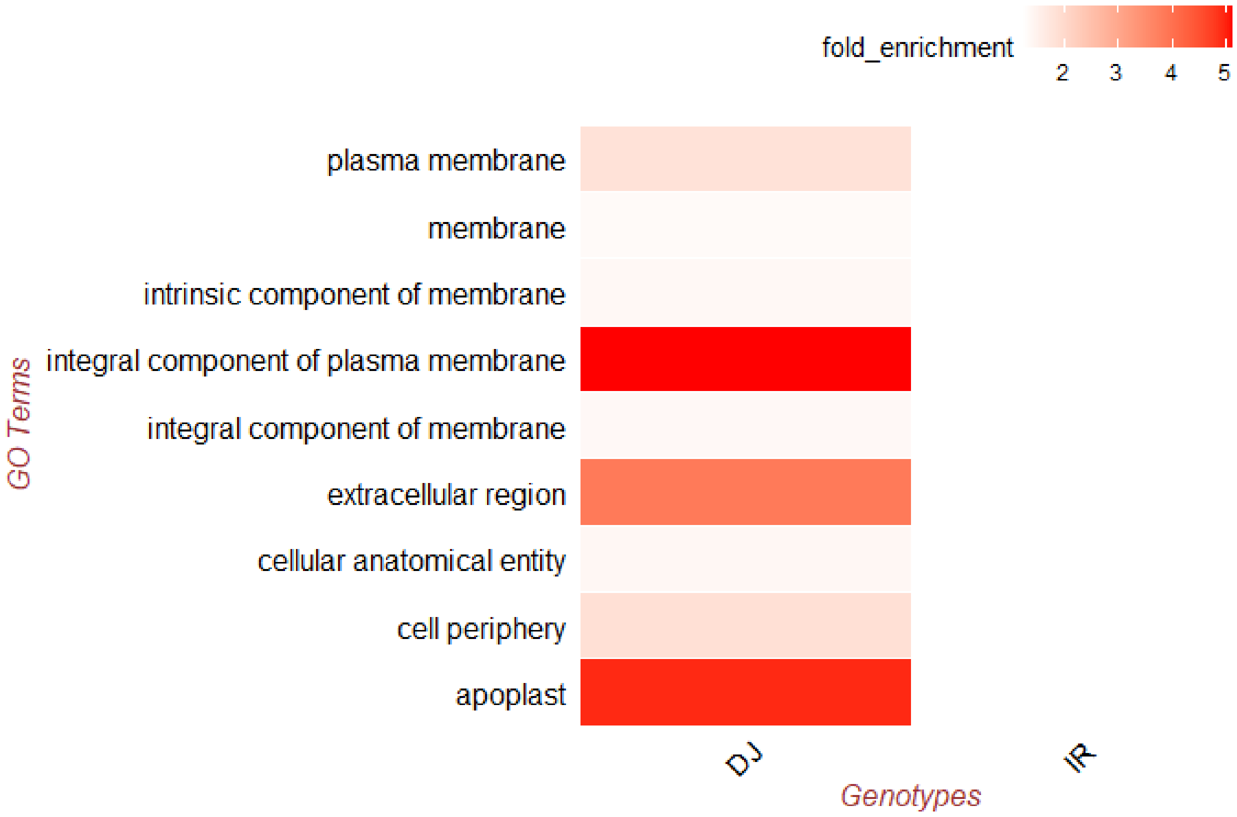


**Supplementary Figure 2.** Cellular Component (GO terms) enriched by the down-regulated genes in DJ123 and IR64 roots under low P treatment.

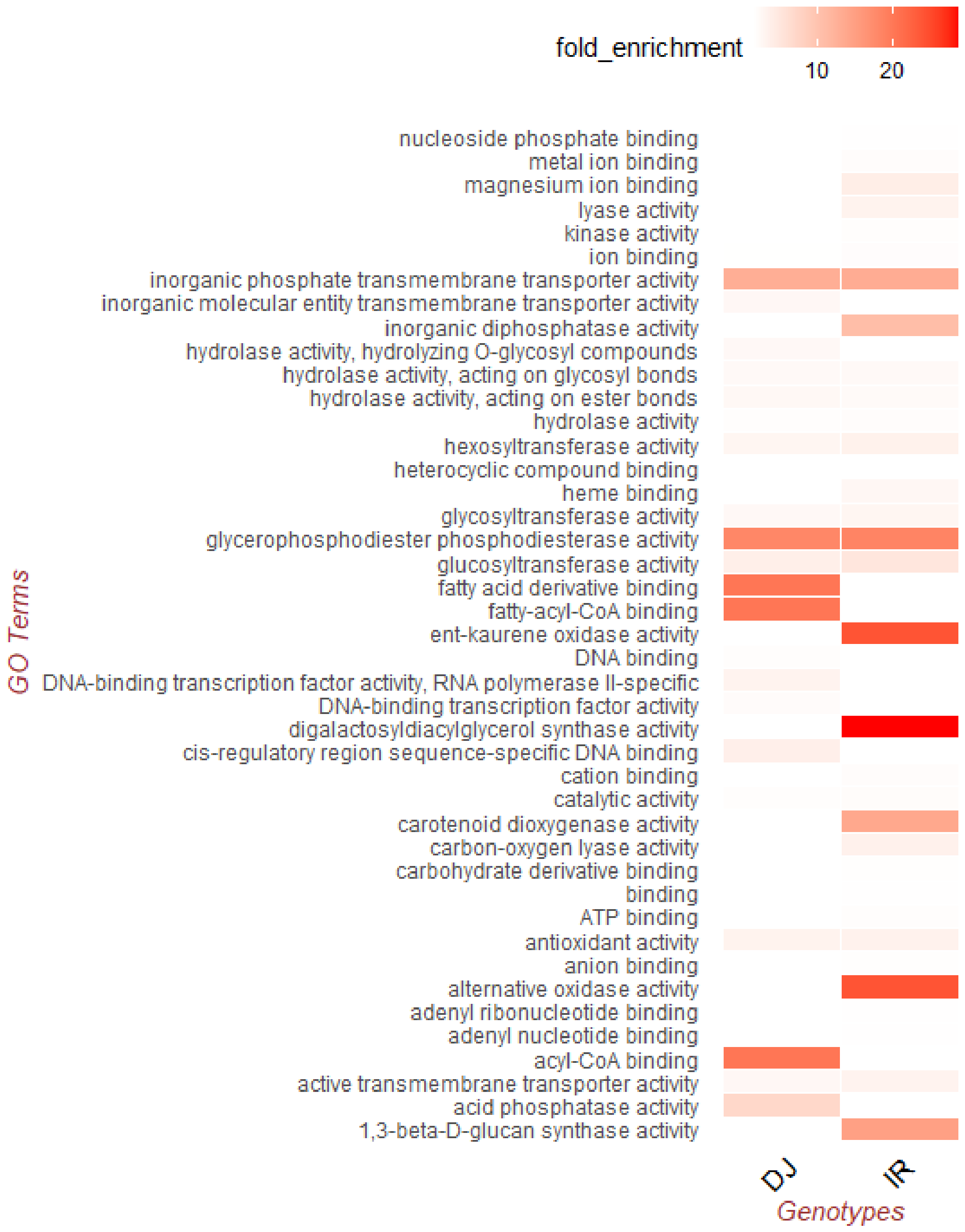


**Supplementary Figure 3.** Molecular Function (GO terms) enriched by the up-regulated genes in DJ123 and IR64 roots under low P treatment (Part1).

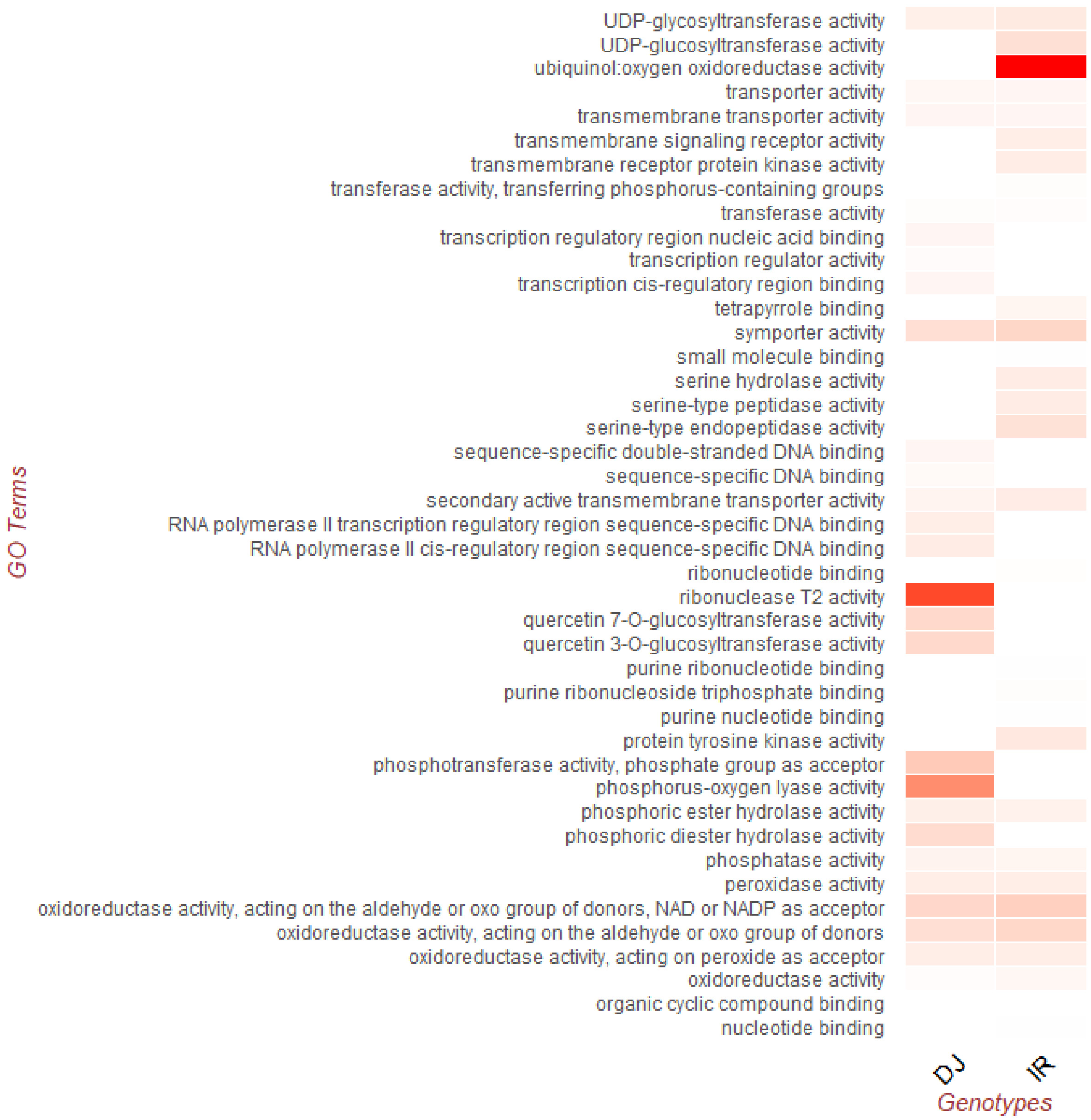


**Supplementary Figure 4.** Molecular Function (GO terms) enriched by the up-regulated genes in DJ123 and IR64 roots under low P treatment (Part2).

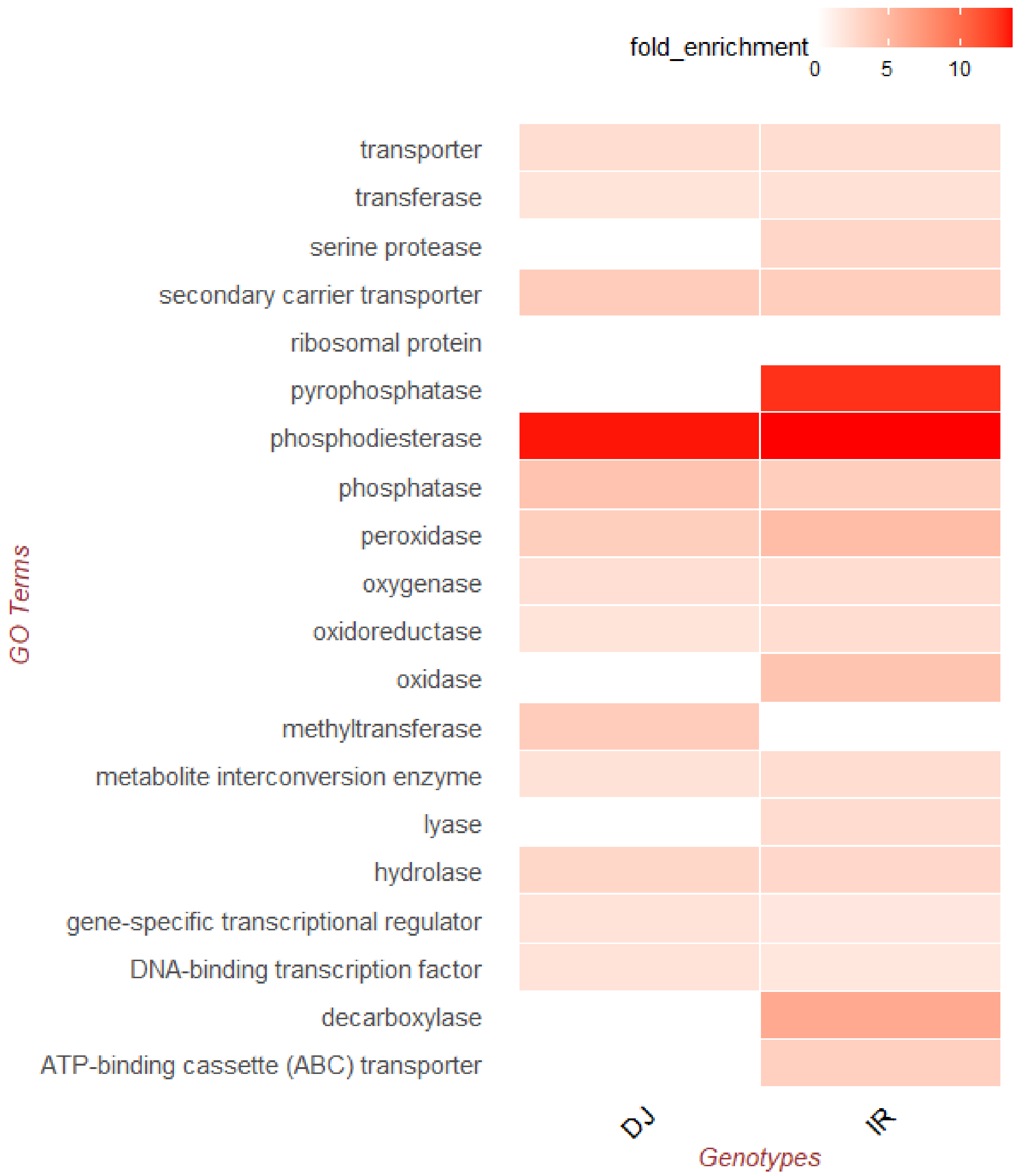


**Supplementary Figure 5.** Protein Class (GO terms) enriched by the up-regulated genes in DJ123 and IR64 roots under low P treatment.

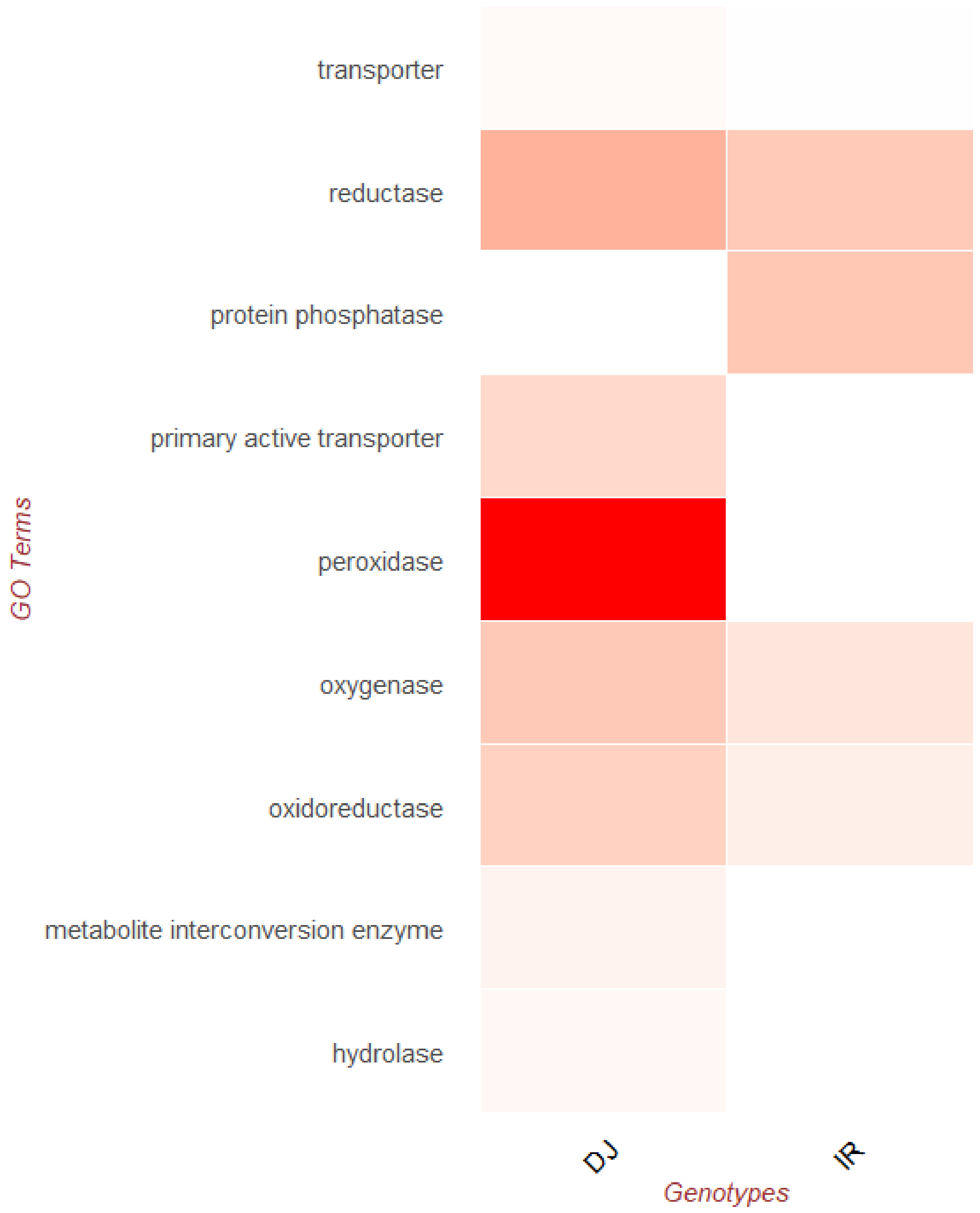


**Supplementary Figure 6.** Protein Class (GO terms) enriched by the down-regulated genes in DJ123 and IR64 roots under low P treatment.
